# Mesowestern Blot: Simultaneous Analysis of Hundreds of Sub-Microliter Lysates

**DOI:** 10.1101/2021.11.07.467614

**Authors:** Cameron O. Zadeh, Jonah R. Huggins, Baylee C. Westbury, William R. Interiano, S. Ashley Phillips, Cemal Erdem, Deepraj Sarmah, William B. Dodd, Wesley O. Meredith, Marc R. Birtwistle

**Affiliations:** Department of Chemical and Biomolecular Engineering, Clemson University, Clemson, SC

## Abstract

Western blotting is a widely-used technique for molecular-weight-resolved analysis of proteins and their post-translational modifications, but has been refractory to affordable scale-up. Here, we report the Mesowestern blot, which uses a 3D-printable gel-casting mold to enable affordable, high-throughput Western blotting with standard sample preparation and small (<1 uL) sample sizes. The casted polyacrylamide gel contains 336, 0.5 uL micropipette-loadable sample wells arranged within a standard microplate footprint. Polyacrylamide % can be altered to change molecular weight resolution range. Proof-of-concept experiments using both infrared-fluorescent molecular weight protein ladder as well as cell lysate (RIPA buffer) demonstrate protein loaded in Mesowestern gels is amenable to the standard Western blotting steps. The main difference between Mesowestern and traditional Western is that semi-dry horizontal instead of immersed vertical gel electrophoresis is used. The linear range of detection is approximately 2 orders of magnitude, with a limit of detection (for β-actin) of around 30 ng of total protein from mammalian cell lysates (~30-3000 cells). Because the gel mold is 3D-printable, users have significant design freedom for custom layouts, and there are few barriers to adoption by the typical cell and molecular biology laboratory already performing Western blots.

## Introduction

The Western blot has been a staple of molecular biology research for decades since its first description in 1979^1^. It uses immersed tank-based polyacrylamide gel electrophoresis to separate proteins by molecular weight, followed by transfer to a nitrocellulose or PVDF membrane, and finally application of antibodies to sensitively detect levels of proteins, post-translational modifications, and even protein complexes^2–4^. Detection modalities include enzyme-mediated generation of colorimetric molecules or light, or direct conjugation of fluorescent molecules to antibodies^5,6^, which when combined with carefully designed experiments, can be quantitative^7,8^. The relatively low cost for traditional Western blotting apparati coupled with ease of use and compatibility with many biological sample types leave Western blotting still widely ingrained in biomedical research as a protein analytic tool, even perhaps the most used technique in protein-related publications in the last 10 years^9^. In fact, the use of western blotting, despite falling “out of fashion” seems stable according to publication metrics^9^.

Despite Western blot usage remaining high, there are notable limitations. Reliance on antibodies for detection is increasingly criticized^10,11^, although separation of proteins by molecular weight is a strong indicator of antibody validity not typically available to other antibody-based technologies—Western blotting is often used as confirmatory assay to bolster support generated by other protein assays. Multiplexing is limited to a handful of analytes per gel, which can be increased slightly by stripping antibodies from the membrane and reprobing with new antibodies^5,12^, cutting the membrane into targeted molecular weight range strips for incubation each with different antibodies^13,14^, or orthogonal detection methods^15,16^. Lastly, traditional Westerns are limited by throughput and sample size; typical gels contain only ~10 wells for analysis of 10 samples simultaneously, and each sample usually requires ~10 ug of total protein content from cell or tissue lysates.

There are several other protein assays that address shortcomings of the Western blot. Reverse phase protein arrays (RPPA) use lysates similar to Western blotting, but greatly increase multiplexing by spotting lysates on chips so that hundreds of antibodies can be used simultaneously^17,18^. However, lysates are not separated by molecular weight, which causes increased stringency for antibody quality; in fact, antibodies are often validated for use in RPPA by Western blot. Luminex xMAP technology offers similar advantages as RPPA^19^. Enzyme-linked immunosorbent assay (ELISA) has been in use even longer than the Western blot, and uses two antibodies, one to capture the analyte from a lysate and the other to detect the captured analyte, with detection modalities similar to Western blots^20,21^. Although ELISA does not separate analytes by molecular weight, the use of two different antibodies for the same target can, in some cases, compensate for specificity issues with one, although obviously the need for two antibodies can be a drawback itself. ELISA enables high-throughput implementation in multi-well plates for simultaneous analysis of hundreds of samples. Mass spectrometry-based proteomics is antibody-free, and can analyze virtually any protein present in a lysate so long as it is ionizable^22–26^. This unmatched multiplexing, however, is hindered by high cost of the apparatus and per sample, difficulty of use for the general biomedical scientist, and large sample sizes often required^5^. Moreover, findings from mass spectrometry experiments often require orthogonal validation with antibody-based techniques such as a Western blot^27^.

There have been advances in Western blotting itself that have improved on the aforementioned limitations. Recent work has enabled single cell Western blotting^28,29^, although this needs a specialized apparatus, so widespread adoption on par with traditional Western blotting is difficult. The Microwestern blot^30–32^ uses a piezoelectric pipetting apparatus to spot small, nL amounts of lysate onto a typical-sized gel, followed by horizontal electrophoresis (as opposed to tank-based), and finally, a gasket system for incubating different parts of the resultant membrane with up to 96 different antibodies. Thus, the Microwestern addresses both throughput and, to some extent, multiplexing limitations, albeit at reduced molecular weight resolution. However, it has not become a widely used technique. The obstacles to adoption center around the piezoelectric pipetting apparatus: (i) it is expensive, difficult to use, and can be mechanically unreliable; (ii) it imposes strict, non-standard sample preparation requirements; (iii) it causes sample loss in tubing dead zones and on the gel, which is flat and does not contain wells. Thus, a Western blotting technique that removes the reliance on piezo-electric pipetting may be of more general use.

Here, we present the Mesowestern blot that, similar to the Microwestern, allows for high-throughput analysis of hundreds of samples in a typical sized gel, but does not require piezoelectric pipetting. To do this, we designed and 3D-printed a gel-casting mold that produces a polyacrylamide gel with 336, 0.5 uL sample wells arranged with 8 rows by 42 columns that is micropipette-loadable. However, the format is flexible because the cast is 3D printed. Proof-of-concept experiments using both infrared-fluorescent molecular weight ladder as well as cell lysates demonstrate that proteins loaded in Mesowestern gels are amenable to the standard Western blotting steps of gel electrophoresis followed by transfer to a membrane for imaging. The main difference from Western blotting is horizontal electrophoresis as opposed to tank-based electrophoresis, but such horizontal apparati are relatively easy to use and not cost-prohibitive. Because the gel mold is 3D printable, users have significant design freedom for custom layouts and we expect that the technique could be easily adopted by any typical cell and molecular biology laboratory already performing Western blots.

## Results

### The Mesowestern Process

The Mesowestern process (Fig. 1A) begins with casting a 1.2 mm thick polyacrylamide gel in the 3D-printed mold (Fig. 2). The mold itself consists of two pieces, the “top” and “bottom”. The top contains the ports in which the unpolymerized gel is loaded, and the bottom contains the impressions of the microwells into which lysates will be loaded after casting. Each microwell negative is roughly a trapezoidal prism of 0.5 mm height, and is slightly longer in one length direction, for a total volume of a little over 0.5 uL. The entire mold has dimensions of a microplate (~9 by 13 cm), and contains eight rows of microwell negatives, with each row containing 42 columns, for a total of 336 potential microwells per gel. Between rows there is ~ 9 mm for proteins to separate, and there is ~2 mm between microwells in the same row.

**Figure 1.**
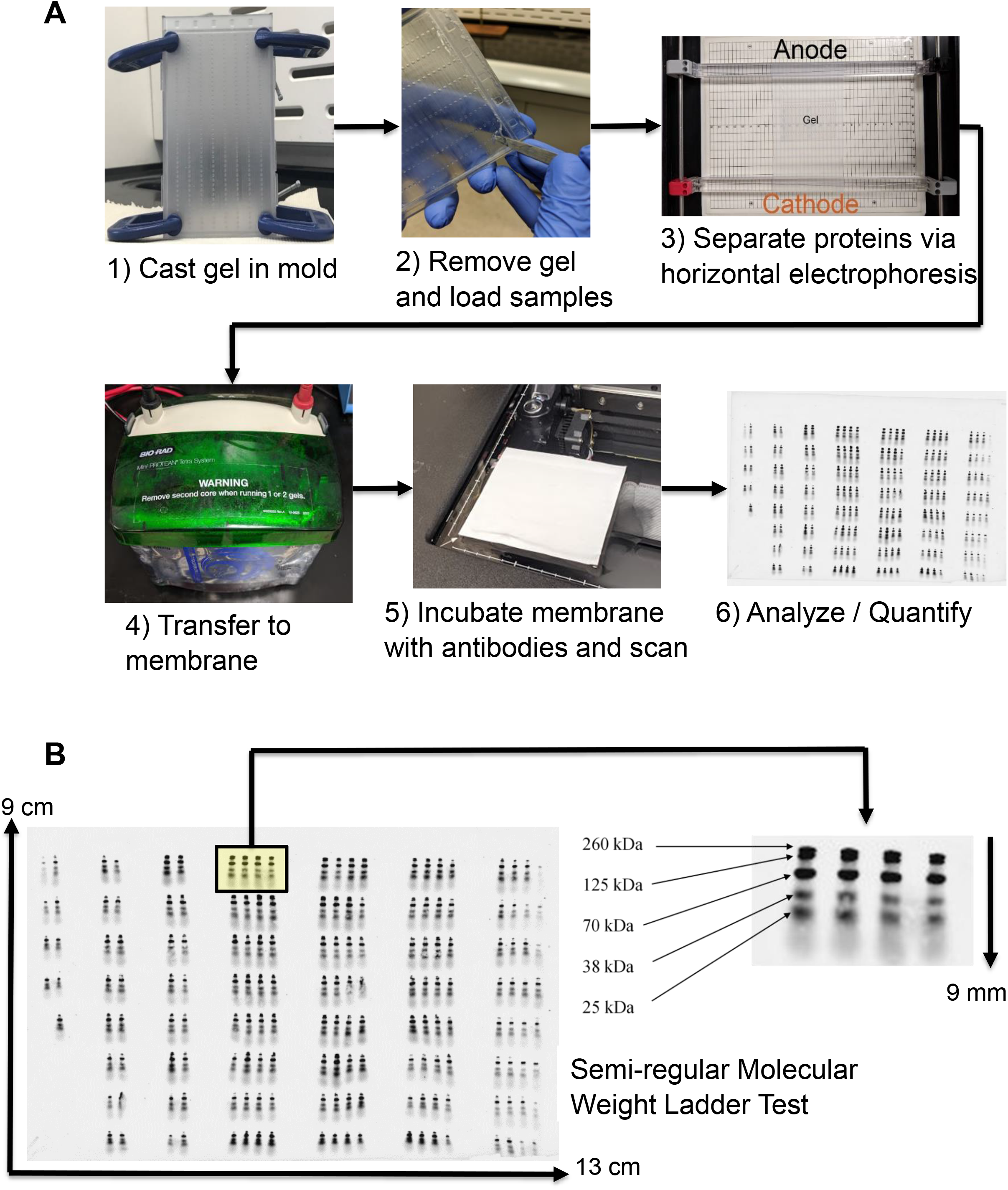
Mesowestern Process and Example Results. **A.** First, the gel is polymerized in the gel casting mold, and after polymerization, it is removed so that samples can be loaded as desired. Electrophoresis takes place on a horizontal apparatus, which is the main difference between Mesowestern and regular Western. After electrophoresis, the Mesowestern and regular Western workflows are identical, with transfer to membrane (tank-based is shown), scanning / visualization, and analysis. **B.** An example Mesowestern membrane where molecular weight ladder was loaded in semi-regular patterns for illustrative purposes. The entire membrane scan is roughly of microplate dimensions, and one “block” of ladder is highlighted. A Mesowestern lane is only approximately 9 mm, but resolves molecular weights between 125 kDa and 25 kDa reasonably well (at 9.5% acrylimide used here).

**Figure 2.**
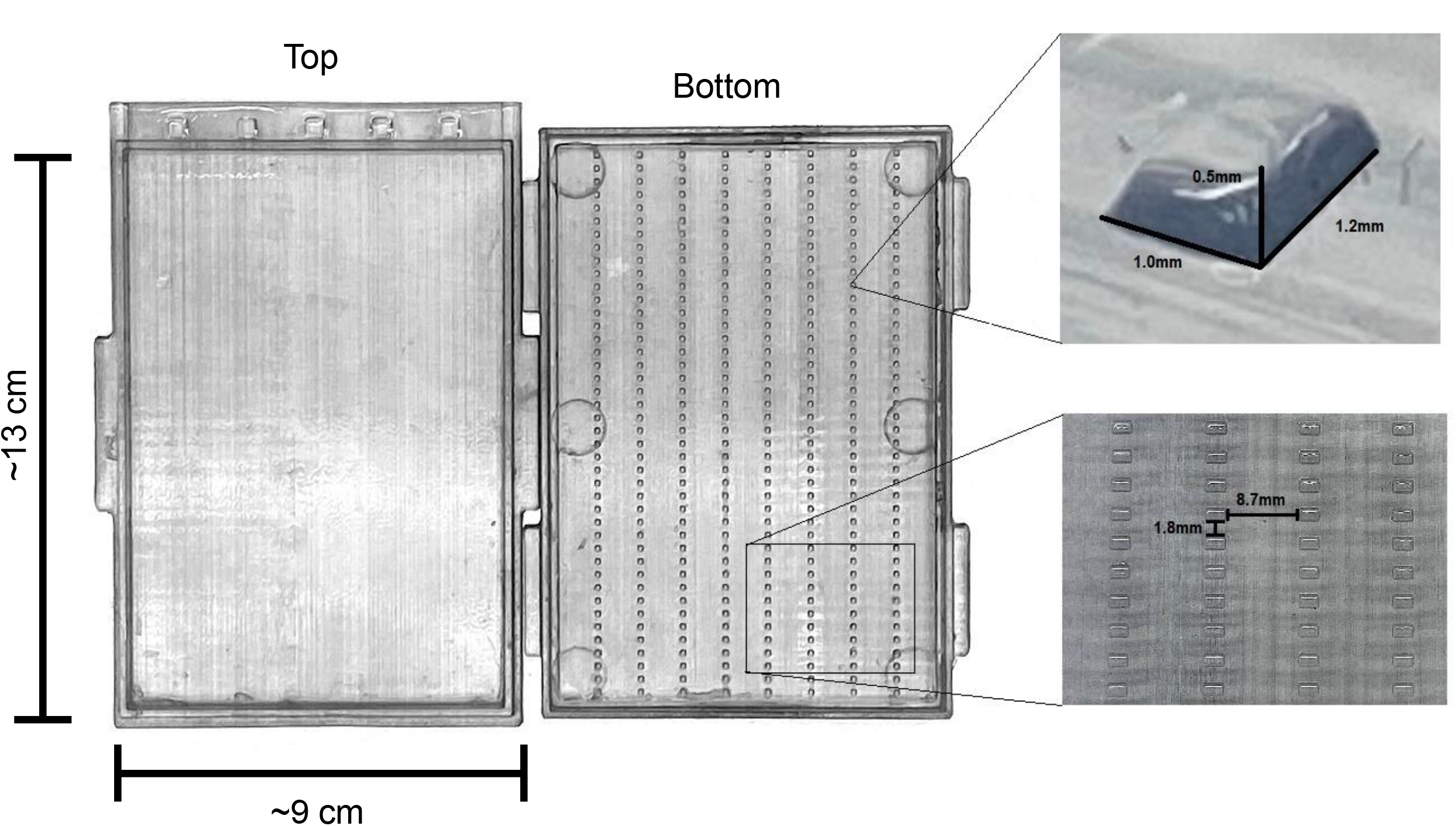
Gel Casting Mold. The mold consists of two pieces which we refer to as “Top” and “Bottom”. The top contains the loading port for unpolymerized gel solution, whereas the bottom contains the raised regions which become wells in the Mesowestern gel. The gel dimensions are approximately 9 cm by 13 cm in width and length, and is 1.2 mm thick. Each well is a rectangle that is 1 mm by 1.2 mm in width and length, and is 0.5 mm deep. Wells are spaced 1.8 mm apart, and have 8.7 mm to run in their lane before the next well is reached.

During casting, the mold stands upright and is held tight by household C-clamps while freshly prepared unpolymerized gel solution (see Methods) is loaded into the casting device from the top, very similar to traditional gel casting between glass plates. After polymerization (~30 minutes), the mold top and bottom are taken apart and the gel can be carefully removed for loading of samples in the microwells via micropipette. After sample loading, horizontal electrophoresis separates proteins by molecular weight. This step is the biggest difference from traditional Western blotting which typically uses immersed tank vertical electrophoresis. Following electrophoresis, the workflow is generally indistinguishable from traditional Western blotting. Tank-based transfer can be employed to move the separated proteins to a nitrocellulose or PVDF membrane, membranes are incubated with antibodies (with block / wash steps), and finally scanned for visualization of bands (we use LICOR infrared fluorescence in this work).

As a simple demonstration, we loaded molecular weight ladder in semi-regular patterns throughout a 9.5% acrylamide Mesowestern gel (Fig. 1B). Although the distance each sample has to run (~ 9 mm) is much smaller than the standard Western blot, and there is no “stacking gel” available, protein separation is reasonably uniform throughout the gel and molecular weight standards within the ladder are distinctly observable between 25 kDa and ~125 kDa, and to some extent at 260 kDa with lesser resolution. Overall, this pipeline establishes a Mesowestern workflow that is highly similar to traditional Western but is much higher throughput with smaller sample sizes.

### Comparison of Western Technologies

After establishing the basic Mesowestern workflow, it is instructive to revisit the similarities and differences between that, the “regular” Western, and the Microwestern^31^ (Fig. 3 and Table 1). At the stage of sample preparation, Mesowestern and Western are identical, whereas Microwestern has multiple differences, such as requiring SDS and DTT in the lysis buffer, sonication steps to clear lysate, and spin column-based sample concentration. These differences limit applicability and throughput of the Microwestern, as well as the downstream assay types that are compatible to measure total protein content.

**Figure 3.**
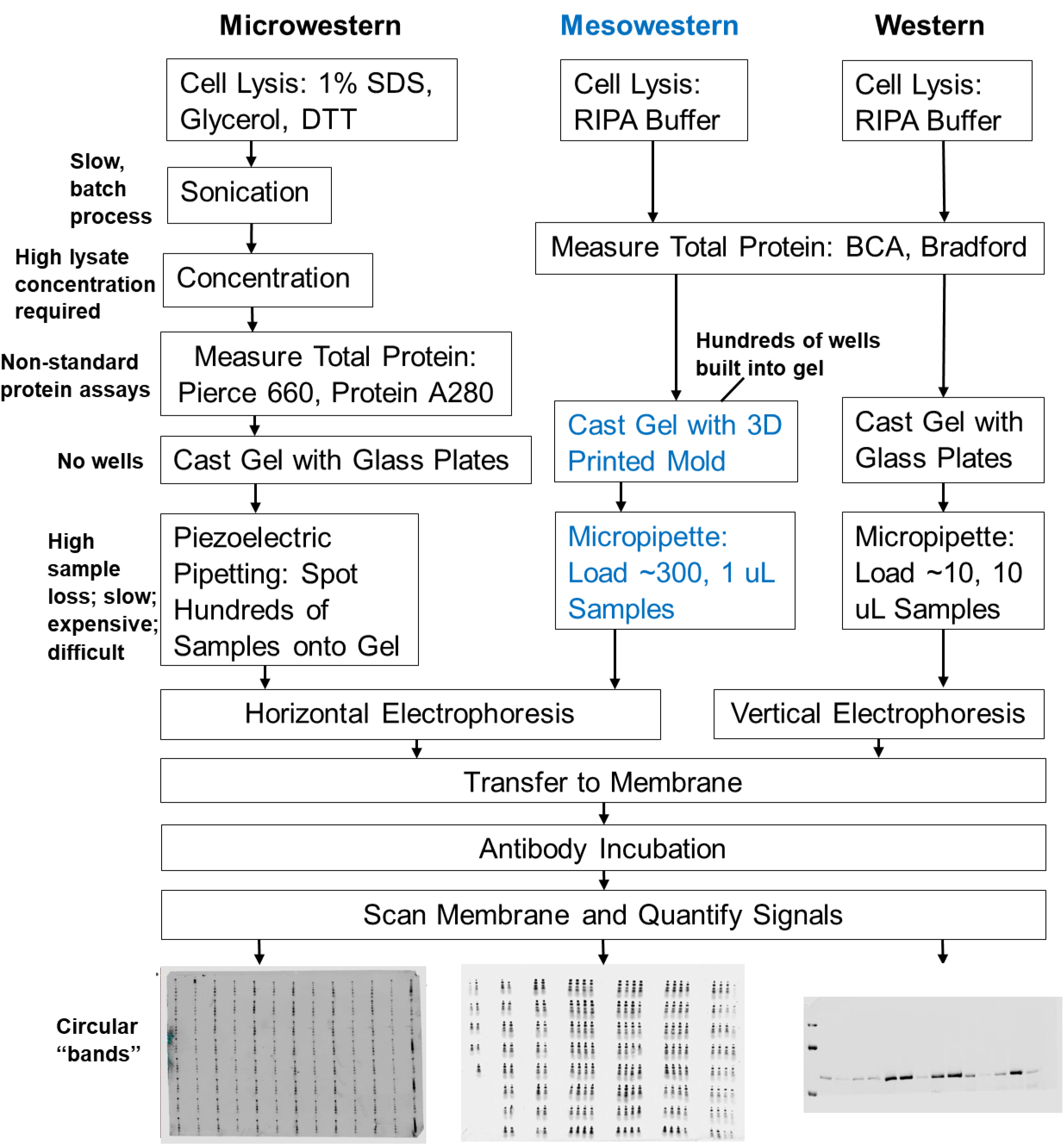
Comparison of Mesowestern to Microwestern and Regular Western. There are significant differences between the Microwestern, the most comparable high throughput western, and the Mesowestern. In terms of processing and workflow, the Mesowestern is very similar to traditional western. The main differences are that the gel is cast with the 3D printed mold, rather than between two glass plates, that much smaller sample volumes are required, and that horizontal electrophoresis is employed. Horizontal electrophoresis is the main point of similarity between Microwestern and Mesowestern. The reliance of Microwestern on a piezoelectric pipetting apparatus creates several upstream problems, including the fact that there are no wells in a Microwestern gel, that lysis buffer is non-standard and lysates must be sonicated and concentrated. After electrophoresis, the workflows of all three processes are the same, but yield quite different images.

**Table 1:** Comparison of each stage of the Western blot, Microwestern Array (MWA) and Mesowestern blot methods.

| Step | Microwestern Array | Mesowestern Blot | Western Blot |
| --- | --- | --- | --- |
| **1. Lysate Preparation** | • ~20 μL per sample volume<br>• Requires sonication<br>• Requires high concentration (~10 mg/mL), necessitating spin columns<br>• Non-standard lysis buffers (e.g. contains SDS) dictates non-standard protein concentration assays | • ~1 μL per sample volume<br>• Standard lysis buffers | • ~10 μL per sample volume<br>• Standard lysis buffers |
| **2. Gel Casting** | • Uses glass plates to cast gels<br>• Requires internal gel support structures (netfix)<br>• Flat gel – no sample wells | • Uses 3D printed mold to cast gels<br>• Hundreds of sample wells | • Uses glass plates to cast gels<br>• Tens of sample wells<br>• Stacking gel used |
| **3. Sample Loading** | • Hundreds of samples can be “spotted” with piezoelectric pipetting<br>• Significant sample loss due to mechanics of piezoelectric pipetting | • Hundreds of ~ 1 μL samples can be loaded with micropipettes | • Tens of ~ 10 μL samples can be loaded with micropipettes |
| **4. Protein Separation** | • Horizontal electrophoresis | • Horizontal electrophoresis | • Vertical (tank) electrophoresis |
| **5. Transfer** | • Wet or semi-dry transfer | • Wet or semi-dry transfer | • Wet or semi-dry transfer |
| **6. Antibody Incubation** | • Incubate different sections or entire membrane with antibodies | • Incubate different sections or entire membrane with antibodies. | • Incubate entire membrane with antibodies. |
| **7. Analysis and Quantification** | • Lower molecular weight resolution due to smaller lanes and circular spotting (no wells) | • Sharper bands than microwestern due to wells. | • Distinct bands with greater molecular weight resolution due to larger lanes / stacking. |

Both Microwestern and Western gels are cast with glass plates, whereas the Mesowestern uses the previously described 3D-printed mold. Importantly, there are no wells in the Microwestern gel, whereas hundreds of small wells are built into the Mesowestern gel cast, and wells are introduced into a Western gel via a low polyacrylamide % stacking gel which promotes subsequent sample focusing. The absence of wells in the Microwestern gel leads to substantial sample loss downstream.

The primary difference with Microwestern is the piezoelectric pipetting-based “spotting” of samples onto the well-less gel, as compared to Mesowestern and Western which both use micropipettes. The hundreds of Mesowestern gel wells hold ~0.5-1 uL of lysate each, whereas the ~10 Western gel wells hold ~10-40 uL of lysate each. While piezoelectric pipetting spots nL-scale samples onto the gel, multiple rounds of spotting are done, and significant sample volume is required to be present in the microwell plate that serves as the sample source, and also in the associated apparatus tubing to ensure robust spotting function. This leads to a substantial amount of lost sample, despite the small amount spotted onto the gel. Lastly, in Microwestern proteins presumably enter the gel through adsorption / diffusion, which leads to circular “bands”, as well as potential sample loss for protein that does not enter the gel via such means.

After samples are loaded, both the Microwestern and Mesowestern use horizontal electrophoresis in a semi-dry setting to separate proteins by molecular weight. Western typically uses immersed tank-based vertical electrophoresis. The latter tends to require less costly equipment and is more readily available, although horizontal electrophoresis apparati are not cost-prohibitive and are simple to operate.

After gel electrophoresis, there are no differences between all three techniques with regards to transfer and preparation for antibody incubation / imaging. The differences manifest with the resultant typical images. Microwestern images are of similar scale to Mesowestern images, but the visualized proteins exhibit circular patterns in Microwestern whereas they are more band-like in Mesowestern. Western, in contrast, has the most sharp bands and largest molecular weight resolution, as stacking gel is used and the proteins are able to migrate significantly longer distances.

We conclude that the Mesowestern offers many benefits compared to Microwestern, namely compatibility with more standard sample preparation workflows, much less sample loss, and elimination of the reliance on piezoelectric pipetting. In comparison to the Western, the Mesowestern offers over 10-fold higher throughput with 10-fold lower sample size requirements.

### Controlling Molecular Weight Resolution by Varying Acrylamide Composition

Given the inherently lower molecular weight resolution of the Mesowestern as compared to the regular Western, we asked whether the acrylamide proportion could be varied in Mesowestern gels to enable more targeted separation of different molecular weight ranges. This is routinely done in regular Westerns. Therefore, we cast gels with 6%, 9.5%, 12% and 18% acrylamide composition (see Methods), and evaluated molecular weight resolution by separation of a ladder standard, relative to the 9.5% case (Fig. 4). In the 9.5% gel, the 160 kDa band has slight mobility, and bands under 30 kDa are not resolvable. In a 6% gel, higher molecular weight proteins should have increased mobility, and the 160 kDa band does have noticeably better gel entry and resolution. In the 12% and 18% gels, lower molecular weight proteins should be more resolved. The 12% gel resolves below 30 kDa better, but still cannot distinguish the 15 and 8 kDa bands. The 18% gel resolves both the 15 kDa and 8 kDa bands. We conclude that varying acrylamide proportion in Mesowestern gels over ranges typically used in regular Western can target a wide range of molecular weights for analysis.

**Figure 4:**
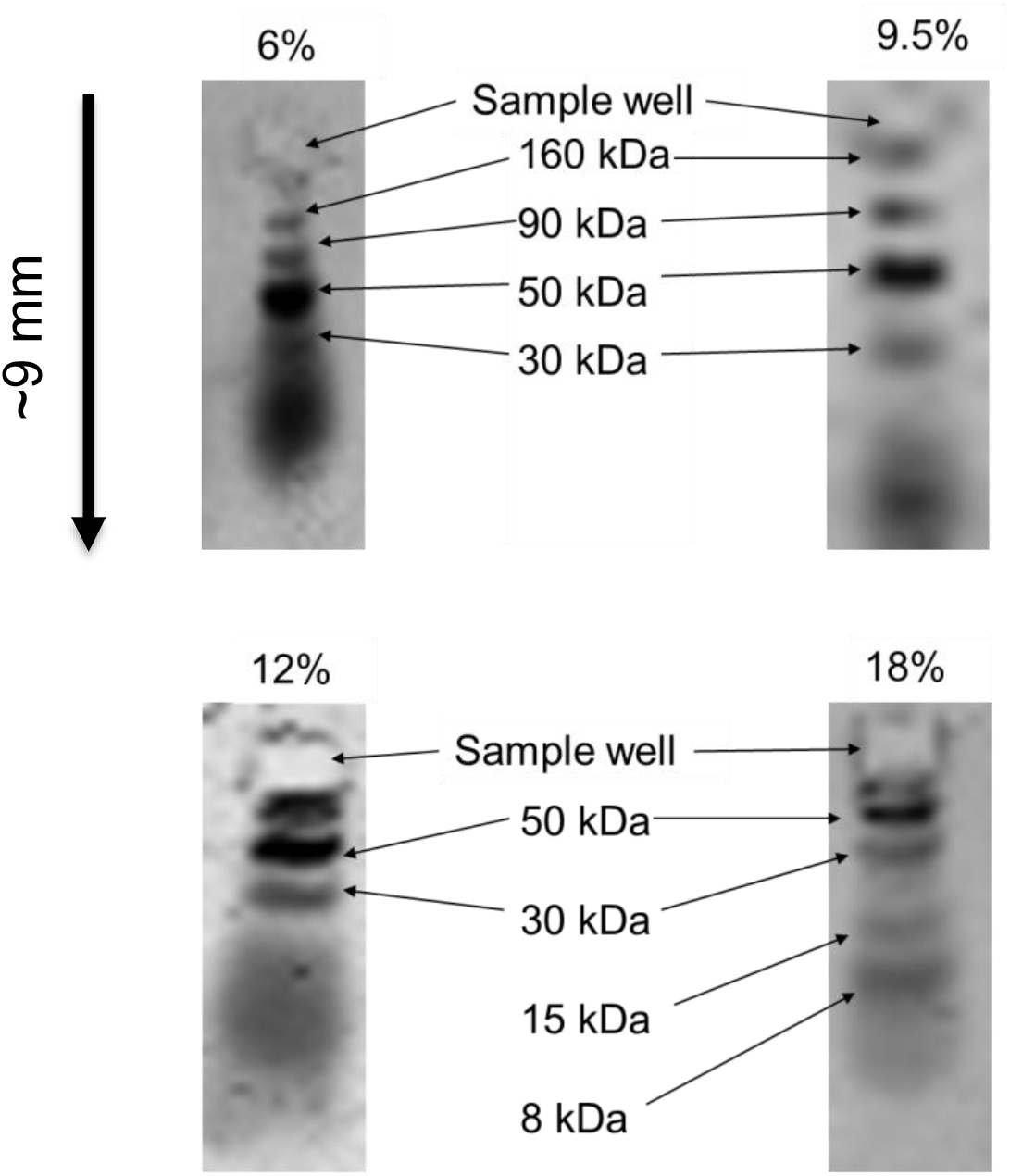
Acrylamide Composition Effects on Molecular Weight Resolution. Representative examples from membranes after transferring molecular weight ladder from four gels at different compositions of acrylamide (denoted by percent) are shown. In general, as expected, higher acrylamide composition resolves lower molecular weights more robustly, at the expense of resolving higher molecular weights. The 9.5% gel gives good general molecular weight resolution.

### Reproducibility Across a Mesowestern Gel

Having established the basic Mesowestern workflow using ladder-based standards, we wanted to evaluate performance using cell lysates and antibodies. The first question we had was related to the reproducibility across a gel. We specifically focused here on a “quarter gel”, which we found often useful, as it still provides high-throughput capability but with reduced labor input. We loaded 0.5 uL of lysate from exponentially growing MCF10A cells into each well of a quarter gel, along with some regularly spaced molecular weight ladder, and then blotted for ⍰-Actin using LICOR infrared fluorescence detection (Fig. 5). The experiment yielded predominantly clear bands at the expected molecular weight (~42 kDa), which a few anomalies not atypical from regular Western. There was noticeable variability in the band intensities, which was found to follow a near normal curve (Fig. 5B). Certainly, heterogeneities in electrophoresis and transfer to membrane could play a role, but we also could not rule out a substantial contribution from small volume manual pipetting. We conclude that the Mesowestern can be used to analyze cell lysates analogously to regular Western. Accordingly, comparing values from direct quantification of bands in different parts of the membrane may be imprecise, and require normalization to additional controls.

**Figure 5:**
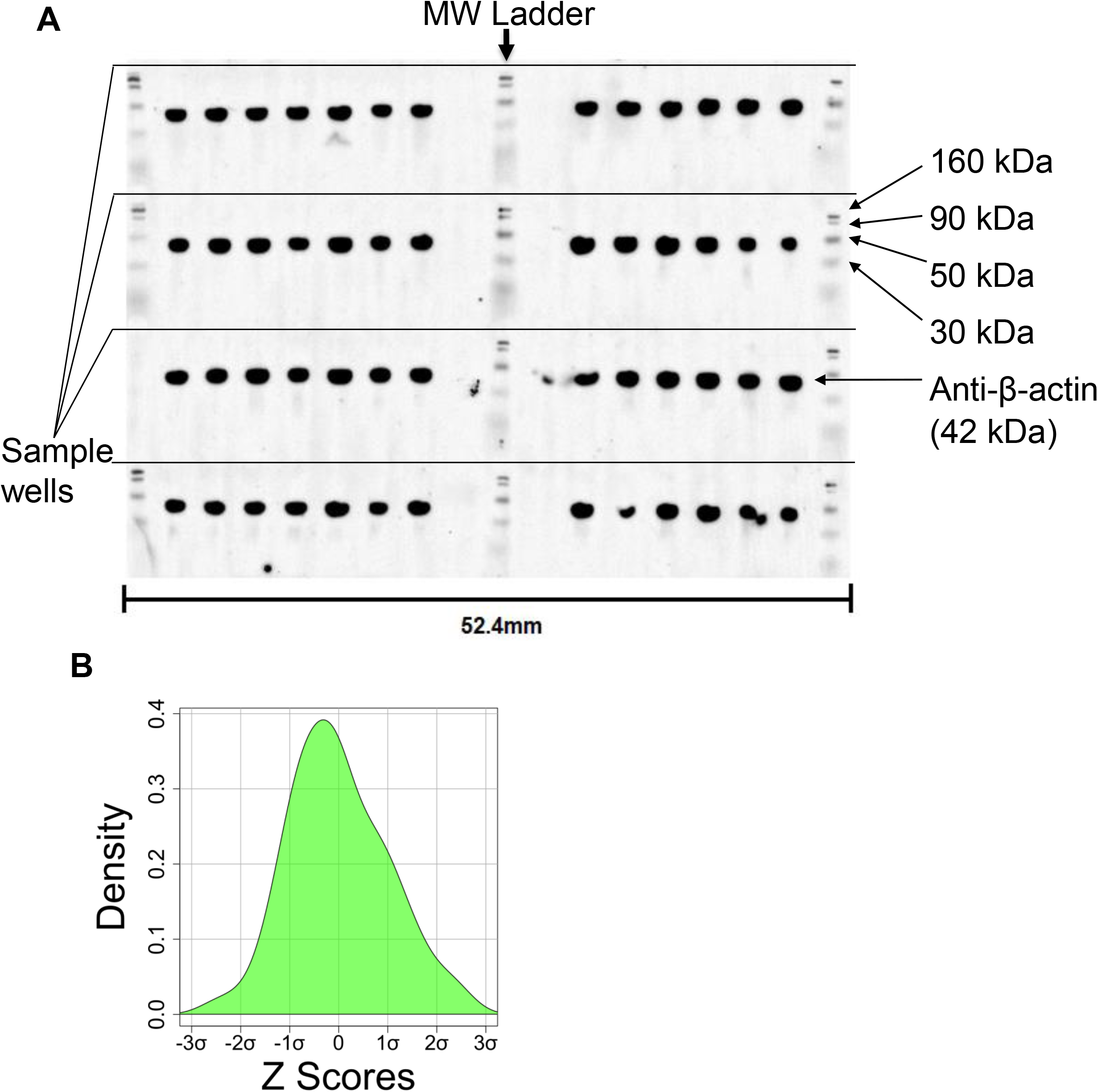
Lysates Analyzed for Reproducibility Across a Quarter Gel. **A.** Exponentially growing MCF10A cells were harvested and lysed as described in Methods. The same lysate sample was loaded into each well of the pictured Mesowestern blot. The total protein concentration of the sample was 4.0 mg/mL and 0.5 μL per well was loaded into the gel. After electrophoresis and transfer, the PVDF membrane was with incubated with anti-β-actin antibodies (1:1,000) and a secondary antibody for detection (see Methods). **B.** We quantified each band in the blot image from A, and analyzed the distribution by z-score analysis as pictured. The distribution is approximately normal, with very little variation outside of 2 standard deviations.

### Dual-Color Imaging Allows Reduction of Variation by Normalization to a Loading Control

One feature of LICOR-based infrared fluorescence approaches is a natural two-color imaging scheme, which in this case, could provide an internal loading control signal for each well with which to improve quantitative comparison from sample to sample. To test this, we again used a quarter gel loaded with cell lysates from exponentially growing MCF10A cells (Fig. 6). As a loading control, we blotted for α-Tubulin, and as an example of a target that may be of interest for quantification, we blotted for doubly phosphorylated ERK1/2 (p-MAPK), a central signal transduction protein. As before, bands were clearly visible at the expected molecular weights (Figs. 6A-B). We quantified these bands for analysis, and found reasonable correlation between their intensities (Fig. 6C-R^2^ = 0.69). To evaluate whether normalizing (i.e. dividing) the p-MAPK signal by the α-Tubulin signal improved the reliability of the p-MAPK signal, we compared the coefficient of variation % (% CV) for each set of values (Fig. 6D). Such normalization reduced the % CV for the p-MAPK signal. We conclude that two-color imaging with internal loading controls may improve quantitative comparability across samples in a Mesowestern gel.

**Figure 6:**
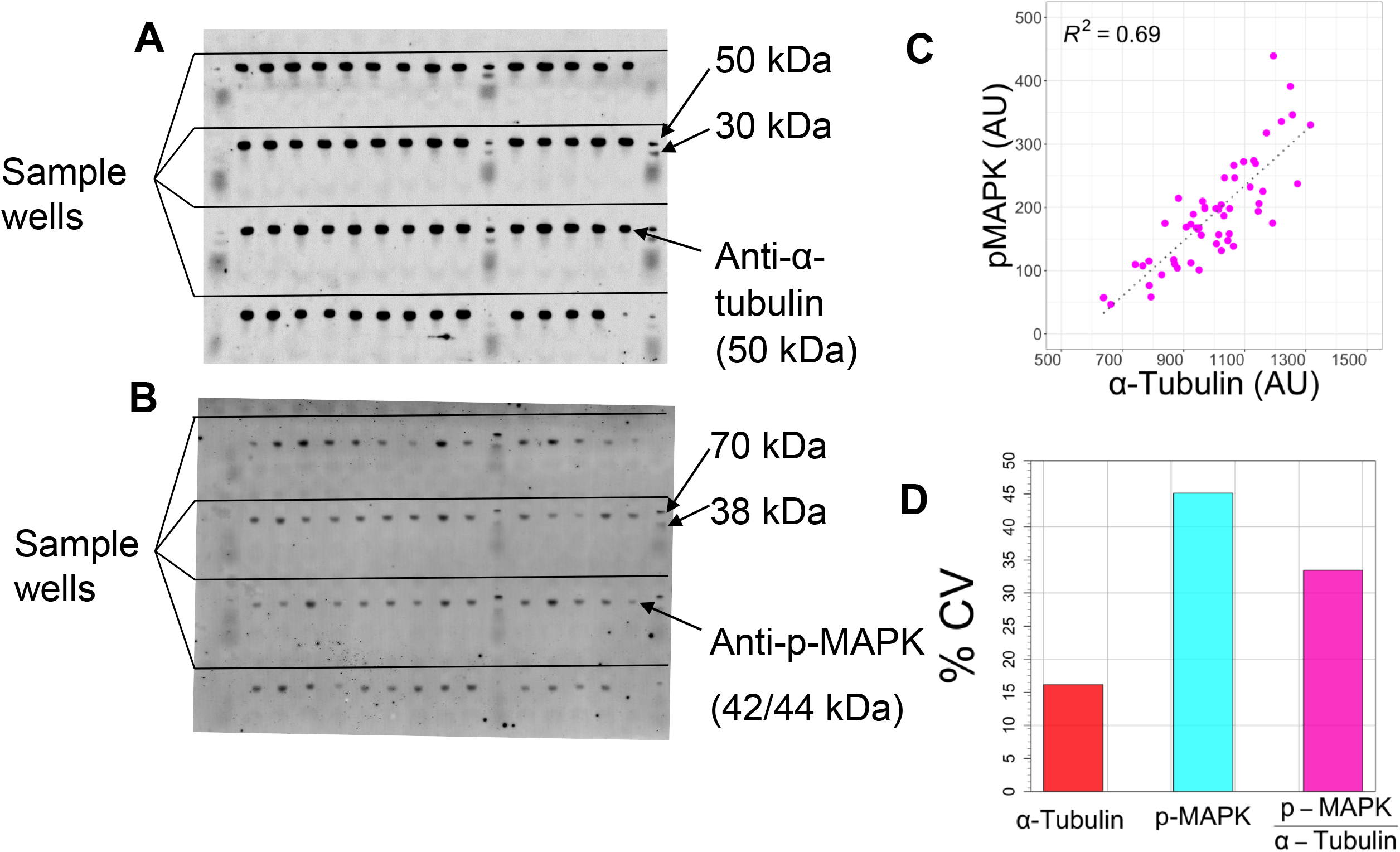
Dual-color Mesowestern Blotting for Loading Control Normalization. **A-B.** Lysate from exponentially growing MCF10A cells was diluted to 2 mg/mL and loaded into a 9.5% gel. After transfer, the membrane was incubated with anti-α-tubulin and anti-p-MAPK antibodies, and then different secondary antibodies for detection of each at different wavelengths. The membrane images depict the same membrane, but at different wavelengths to detect **A.** α-tubulin and **B.** p-MAPK separately. **C.** Quantified bands were plotted to examine the expected correlation between the two signals from the same lysate. **D.** The variation across the gel of the independent signals, and of the p-MAPK signal normalized by the α-tubulin loading control. Dividing by loading control signal improves the %CV.

### Linear Range and Limit of Detection

Because the Mesowestern deals with much smaller sample size than the regular Western, we wanted to determine what the linear range and limits of detection were. While the answer to this question will invariably be dependent on the epitope of interest, its abundance in the cell lysate, and the antibody being used, we started by investigating this for a highly expressed protein, β-Actin. Specifically, we performed a 7-point, 2-fold serial dilution of lysate from exponentially growing MCF10A cells, and replicated this dilution curve 8 times on one Mesowestern quarter gel (Fig. 7A). In many cases, the very last sample, corresponding to ~30 ng of total protein from the lysate, was detectable. However, this was not always the case, again highlighting the variability that may be present across the gel / membrane on a sample-by-sample basis. To evaluate linearity, we quantified each observable band, and then plotted this intensity versus the known amount of lysate loaded (total protein content), which yielded R^2^ = 0.87 (Fig. 7B). We conclude that quite small amounts of protein may be detected by Mesowestern, and at least in the case of MCF10A lysates and β-Actin, as little as 30 ng of total protein from lysate can be detected. It is instructive to reiterate that this is not 30 ng of pure β-Actin, but rather 30 ng from all the protein in the MCF10A cell lysate, of which only a small fraction is β-Actin. This signal is approximately linear at least for 2 orders of magnitude.

**Figure 7:**
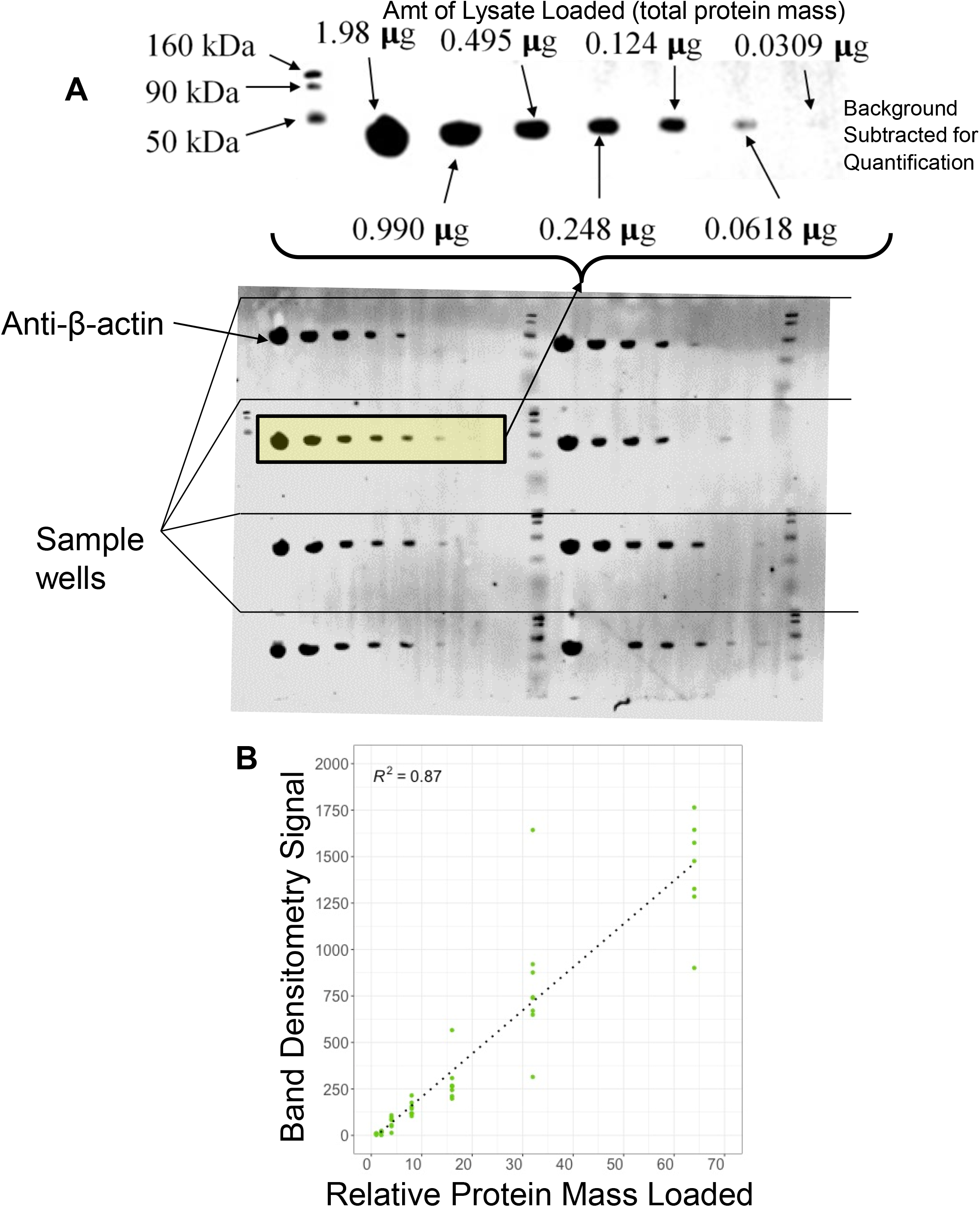
Linear Range and Limit of Detection Experiments. **A.** Cell lysate from exponentially growing MCF10A cells was prepared at a range of protein concentrations (twofold serial dilution) and subjected to Mesowestern analysis as indicated. The membrane was incubated with anti-β-actin antibodies and a secondary antibody for detection. **B**. The signal derived from image analysis of each band was plotted versus the known amount of total protein mass loaded. Good correlation is observed and these experiments lie within the linear range.

## Discussion

Here we have described the development and first functional testing of a high-throughput, small sample size Western blotting protocol called Mesowestern. As compared to a Western blot, it enables at least 10-fold greater sample throughput with at least 1/10 the amount of lysate per sample, and only requires horizontal as opposed to the traditionally-employed vertical electrophoresis. As compared to the Microwestern, it eliminates the need for piezoelectric pipetting by using a 3D-printed gel mold that makes micropipette-loadable gels. We have demonstrated that molecular weight resolution of a Mesowestern gel can be adjusted in expected ways by changing gel acrylamide composition. We have explored the limits of detection, linear range, and quantitative ability of the Mesowestern, which was found to be similar to regular Western. Overall, the Mesowestern is a promising technology that could be readily adopted by molecular biology labs having interest in more high-throughput Western blotting with small samples sizes.

Although we have used here a single mold design, the layout can be quite easily modified as users may desire for their particular applications. Access to the necessary 3D printing facilities is relatively common and the printing itself is fast and inexpensive, so we anticipate that custom molds will be easy to implement. For example, some users may want fewer wells, but to be able to load more sample volume per well. Others may want more separation space available to each well. Yet others may wish to make an even larger gel, much larger than a microwell plate footprint (compatible with the downstream horizontal electrophoresis). All such variations are straightforward and possible depending on the needs that arise.

One difference between the Mesowestern and regular Western is a “stacking” portion in a regular Western gel^33^. The stacking gel has a low acrylamide composition (~5%) with the purpose of allowing all the proteins from the cell lysate to easily enter the subsequent “resolving” gel at the same time, which presumably allows for crisper bands and thus better molecular weight resolution. In the Mesowestern, there is only a resolving gel. In our applications so far, we indeed have noted that the resulting bands do not have the “crispness” similar to regular Western blots, although they were certainly identifiable at the expected molecular weight ranges. However, given the fact that Mesowestern inherently has lower molecular weight resolution due to less distance for proteins to migrate, future innovations incorporating stacking portions would be a welcome development. This is quite challenging, however, as the unpolymerized gel is loaded from a single entry port, making it difficult to isolate spatial regions where wells reside and stacking gels would be appropriate. In that regard, the ability to cast gradient Mesowestern gels would also be welcome, but similarly challenging.

Lastly, although the Mesowestern makes significant advances with regards to throughput and sample size, we have not demonstrated here the multiplexing capabilities offered by Microwestern. The Microwestern achieves high multiplexing (e.g. 96 antibody pairs are once) by placing the resultant membrane in a microwell plate-sized gasketing apparatus, which allows different antibodies on different parts of the membrane. There is no barrier to applying such an approach to the Mesowestern, so we expect that similar multiplexing can be done, albeit of course at the cost of being less high throughput, since wells must contain repeated patterns of the same lysates to be then blotted by different antibodies.

In conclusion, we have demonstrated here a new approach to Western blotting called the Mesowestern that increases throughput greater than 10-fold with greatly reduced sample size requirements. Most notably, the Mesowestern is straightforward to implement in typical cell and molecular biology labs. While the Western blot may be viewed by some as “old” and “irrelevant”, it does in fact remain as one of the most widely used assays in biomedical science^9^, and this is unlikely to change due to its popular use as a sensitive and specific confirmatory assay modality. Thus, improvements to Western such as the Mesowestern we developed here are still expected to have widespread impact.

## Methods

### Printing the Mold

Molds were printed in the Clemson Additive Manufacturing Lab with the Stratasys Connex 350 and Veroclear (Stratasys, OBJ-03271-RGD810) as the material. Following printing, self-forming valve packing (Danco, #80794) was inlayed into the outer edge of the well perforation unit (bottom). Schematic files are available upon collaborative request.

### Casting a Gel

Gel casting was completed through a process of silanization of surfaces coming into contact with the gel, clamping to ensure a tight leak-proof fit, and serological pipetting of unpolymerized solution into the mold. Briefly, 2.5% v/v silane solution was prepared by combining 1.25 mL dichlorodimethylsilane (SigmaAldrich, #40140) and 48.75 mL 100% ethanol (Fisher, #04-355-22) in a 50 mL conical tube (Fisher, 14-432-22). We then applied 250 uL of the silane solution to the interior surfaces of both the top and bottom mold pieces, gently spread it across the surface by rocking, and wicked excess with a kimwipe. After assembling the top and bottom pieces together, four C-clamps (Irwin #1901235) were tightened onto the assembly at the designated locations (indented circles). At this point, the assembly is ready for loading.

A 9.5% gel solution was prepared by combining 47.5 mL 30% Bis/Acrylamide solution 29:1 (BIO-RAD, #161-0156) with 41mL MilliQ Water, 30mL Glycerol (Sigma, #G5516-500mL), 30mL 5x Tris-Acetate Buffer (recipe as follows), and 1.5mL 10% Sodium Dodecyl Sulfate (SDS) (Fisher, #BP2436) together, respectively. Preparation of the 5x Tris Acetate buffer was completed by dissolving 145.4 g Tris base (BIO-RAD, 161-0719) in 700 mL of MilliQ water (pH expected between 11.0 and 11.4). The pH was adjusted by adding 65 mL glacial acetic acid (Sigma-Aldrich, #320099) and letting the solution sit overnight. Then, 0.5 mL glacial acetic acid was pipetted into the solution and allowed to sit for an hour at room temperature. This was repeated until the solution reached pH 6.9. Finally, the volume of the solution was brought up to 1L with MilliQ water and stored at 4°C.

Polymerizing gel solution was made by combining 15 mL of 9.5% gel solution with 133 uL of 10% Ammonium Persulfate Solution (APS) and 13.3uL TEMED (BIO-RAD #161-0700) into a beaker under a fume hood. The 10% APS solution was prepared by dissolving 0.2g ammonium persulfate (BIO-RAD #161-0800) into 2mL MilliQ water. Quickly after preparation, 15mL of gel solution was dispensed by serological pipetting into the mold assembly via the loading port (Fig. 1). The assembly was kept still under the fume hood for 30 minutes at room temperature to achieve full polymerization.

To remove the gel from the mold, first the C-clamps were removed. Then the top and bottom mold pieces were carefully separated using a gel releaser (BIORAD, #165330) on the lateral protrusions, followed by carefully moving the releaser around the internal face of the top. After splitting the top and bottom pieces, the gel is removed by inverting the mold so that the gel is facing thick blotter paper (BIORAD #1703958) that is presoaked in running buffer (see below). The blotter paper was approximately 5 cm larger than the gel on the top and bottom, and about 1 cm larger than the gel on each side. The gel is slowly peeled away from a corner using the gel releaser until gravity facilitates the remaining gel to gently fall onto the soaked blotter paper support. The gel can be used immediately or be stored in a sealed bag at 4°C for several months (at least).

### Cell Culture

MCF10A cells (from LINCS Consortium and STR verified internally) are cultured in DMEM/F12 (Gibco #11330032) medium containing 5% (vol/vol) horse serum (Gibco #16050122), 20ng/mL EGF (PeproTech #AF-100-15), 0.5 mg/mL hydrocortisone (Sigma #H-0888), 10μg/mL insulin (Sigma #I-1882), 100ng/mL cholera toxin (Sigma #C-8052), and 2mM L-Glutamine (Corning #25-005-CI). The cells are kept at 37°C in 5% CO_2_ in a humidified incubator. To maintain subconfluency, the cells are passaged every 2-3 days, washing once with PBS (phosphate buffered saline), lifting with 0.25% trypsin (Corning #25-053-CI), and reseeding in full growth media.

### Lysate and Sample Preparation

Cells growing in full growth media were collected, counted, and seeded (150,000 cells/well) in tissue culture treated six well plates (Corning # 08-772-1B). The cells are kept at 37°C in 5% CO_2_ in a humidified incubator for ~48 hours. The plates were removed from the incubator and media in the wells aspirated. The wells were washed with icecold PBS once and placed on ice. Freshly prepared, ice-cold RIPA buffer (110 μL, 50mM Tris, pH 7-8 (Acros Organics #14050-0010), 150 mM NaCl (Fluka #71383), 0.1 % SDS (Fisher #46040CI), 0.5% sodium deoxycholate (Alfa Aesar, J62288), 1% Triton-X-100 (Fisher, BP151) with protease & phosphatase inhibitors (1μg/mL aprotinin (MP Biomedicals #0219115801), 1μg/mL leupeptin (MP Biochemicals #0215155301), 1μg/mL pepstatin A (MP Biochemicals #0219536801), 10 mM β-glycerophosphate (Santa Cruz Biotechnology #sc203323), and 1mM sodium orthovanadate (Sigma # S6508)) are added into each well. The plates are kept on a rocker (slow) in the cold room for 15 minutes. Then, the lysates are scraped off with a cell scraper (Stellar Scientific TC-CS-25) and 100 μL lysate from each well is transferred into labeled Eppendorf tubes on ice. Each tube is vortexed three times for ~ 30 seconds to homogenize cell debris. Next, the tubes are centrifuged at 4°C for 15 minutes at ~21,000 x g (max speed). Finally, 80 μL of the supernatant from each tube was transferred into a new Eppendorf tube, being careful not to disturb the debris pellet. Lysates were stored at −20°C for short-term storage and transferred to −80°C for long-term storage.

### Protein quantification

Total protein quantification of lysates was done using BCA-Pierce 660 Assay (Thermo Scientific #23225) and BSA stock (Thermo Scientific #23209) is used as reference according to the manufacturer’s protocol. In short, 10 μL of lysate sample or BSA standards are loaded into 96-well plates (Corning #3370), in triplicate. Then, 150 μL BCA Protein Assay Reagent is loaded into each non-empty well. The plate is covered with the lid and incubated at room temperature for 5 minutes. The absorbance readings at 660 nm are obtained in a plate reader (BioTek #Epoch2). The average of blank wells is subtracted from each reading to calculate blank-corrected averages for each condition. The standard curve is fitted by a polynomial using blank-corrected mean values of each standard condition versus its BSA concentration. The protein concentration in each sample is calculated using the standard curve formula.

### Sample Preparation

Lysate stocks are thawed on ice (if applicable). Then, 5X Sample Buffer was prepared (5mL of Glycerol (Sigma #G5516), 0.5mL 10% SDS (Fisher #BP 2436), 0.01g Bromophenol blue (Calbiochem #2830), 2.1mL 5x Tris-Acetate Buffer (as above), 0.5mL Betamercaptoethanol (Sigma #M6250), then total volume brought to 10mL with MilliQ water). This was mixed with lysates in a 1:4 (v/v) ratio. Next, the tubes were heated at 95°C for 5min in a dry heating block, and then briefly spun in benchtop microcentrifuge before loading (below).

### Loading the Gel

Following release of the gel onto the soaked blotter paper, the assembly was placed down on a flat surface with the wells facing up. If folds and stretching of the gel are evident, light rolling was used to flatten. A p2 micropipette with 10 uL tips was used to load 0.5 uL of prepared lysates and/or molecular weight ladder (LI-COR, 928-60000) into wells as desired. We have found that wells less than 2 mm away from the gel boundaries may be subject to inconsistent electrophoresis and transfer, and therefore avoid them when possible. Care was taken not to adjust the gel on the blotter paper after any loading, and also to transport the gel with a spatula support underneath.

### Horizontal Electrophoresis

Horizontal electrophoresis was carried out using the Flatbed Professional (Gel Company Store, FC-EDCProf-2836). The apparatus was maintained at 10°C during electrophoresis. First, ~10 mL of cooled running buffer was poured onto the center of the apparatus, followed by transfer of the blotter paper / loaded gel by spatula onto this buffer. Running buffer was made by combining 20 mL of 5x Tris-Acetate buffer (see above) with 29.5 mL MilliQ water and 0.5 mL 10% SDS. The gel should be oriented to have the cathode (red bar) at the bottom, where the proteins will migrate towards. Additionally, the wells should be aligned with the apparatus gridlines, and excess running buffer should be wiped up with no buffer accumulated outside of the blotter paper. Then, the anode and cathode wires were placed over the blotter paper, about 3 cm from the gel. Finally, the glass plate is placed on top of the anode and cathode and the lid is closed. Electrophoresis is conducted at 100 Volts for ~ 2 hours, although each run should be individually monitored. Samples should be visible as blue dots in the gel after ~30 minutes, and ideally, the run should be stopped when it reaches the top edge of the next well. After 30 minutes, we paused the run, lifted the blotter paper and gel with a spatula, and rehydrated by placing another 10 mL of cool running buffer as previously.

### Transfer to Membrane

Transfer buffer was prepared by first making 10x Tris-Glycine Buffer (600 mL of MilliQ water with 30.3 g Tris base (BIO-RAD #161-0719) and 144 g Glycine (VWR #0167), then MilliQ water added to a final volume of 800 mL). Transfer buffer (~2L, 1x) is made by taking 160 mL of 10x Tris-Glycine buffer, adding MilliQ water up to a final volume of 1600 mL, and finally, adding 400 mL of methanol (Fisher #A412-in a fume hood). Transfer buffer is stored at 4°C.

For quarter gels, we used a Mini Trans-Blot Cell (BIO-RAD, 1703930), and for full gels, we used a Criterion Blotter (BIO-RAD, 1704070). We have successfully used both nitrocellulose (GE, 10600002) and PVDF (BIO-RAD, 1620264) membranes for Mesowestern. In our experience, low fluorescence PVDF membranes tend to provide better signal to noise due to their increased ability to bind low abundance proteins, although answering such questions definitively was not the purpose of this manuscript. For PVDF, the membrane was pre-wet with methanol prior to subsequent use and never allowed to dry out.

To prepare the gel and membrane for transfer, cold transfer buffer was poured into a pyrex dish to a depth of ~ 3 cm. Blotter paper, cut to the size of the transfer cassette but larger than the gel, was placed into the pyrex dish to soak. After soaking, the blotter paper was placed on a clean, flat bench top. Then, the gel was allowed to soak in the same transfer buffer for ~ 15 minutes, making sure to keep track of which side of the gel has the well indentations. The gel was then placed onto the soaked blotter paper, with the wells face down on the paper. A spatula was always used to transport the gel. The gel was then gently rolled flat and air pockets removed using a roller (BIO-RAD, 1651279). The membrane was cut to the same size as the gel, being careful never to touch the membrane except with clean tools. After wetting with methanol (if PVDF is used), the membrane was then placed to soak in transfer buffer. Forceps were used to gently place the membrane onto the gel. If the membrane is not aligned, we did not move it, rather, we got a new membrane. Then, the membrane was rolled as previously. A second piece of transfer buffer-soaked blotter paper was then placed on top of the membrane in line with the first piece of blotter paper, and rolled as previously. Finally, a spatula was used to lift the “sandwich” onto a fiber pad (BIORAD, 1703933), and another fiber pad was placed on top. This fiber pad-surrounded sandwich was moved to the transfer cassette, making sure that the side of the sandwich closest to the membrane was on the clear/positive side of the cassette (BIORAD, 1703931). This also means that the side of the sandwich closest to the gel is on the black/negative side of the cassette. The cassette was then placed into the transfer apparatus (negative to negative/black to black, positive to positive/clear to red). If desired, a second sandwich can be made.

With the cassettes in the transfer apparatus, cold transfer buffer was added until it reached the indicated volume line. The apparatus was moved to a 4°C room, and then transfer was carried out at 30 Volts for 16 hours (usually overnight). After the transfer, the membrane was removed with clean forceps, and was placed in a clean incubation box (Li-Cor, 929-97201), with the side of the membrane that was in contact with the gel facing up.

### Antibody Incubation

First, TBS and TBST buffers were prepared. Briefly, 10X TBS was made by dissolving 24 g Tris Base (BIO-RAD #161-0719) and 88 g NaCl (CAS 7647-14-5) in 1L MilliQ water. The pH was monitored with continuous magnetic stirring, while adding HCl dropwise to bring the pH to 7.6. To make 1X TBS, 50 mL of 10X TBS was added to 450 mL MilliQ water, and stored at 4°C (stable for several months). To make 1X TBST, 2.5 mL of 10% Tween 20 (BIO-RAD #161-0781) was added to 500 mL of 1X TBS, and similarly stored at 4°C.

All membrane incubations are done in the dark (sealed black box or covered in aluminum foil). The membrane was incubated first in ~ 20 mL of blocking buffer (1 g BSA (Fisher, BP1600) in 20 mL of 1X TBS) for at least 30 minutes at room temperature with gentle rocking. Blocking buffer can be reused and stored at −20°C. After blocking, blocking buffer was removed, and the membrane was directly incubated with primary antibody solution (10 mL blocking buffer, 50 uL of 10% Tween 20, v/v dilution of primary antibody to desired working concentration) for at least 2 hours at room temperature or overnight at 4°C, all with gentle rocking. The antibody solution can be saved at −20°C and reused. After primary antibody incubation, the membrane is washed with ~10 mL of 1X TBST three times, 5 min for nitrocellulose, and four times, 15 min for PVDF. After washing, secondary antibody solution (10 mL 1X TBST with 1:20,000 v/v, see below) was added to the membrane and incubated with gentle rocking for 1 hour at room temperature. After incubation, the secondary antibody solution was discarded, and the membrane washed as previously with 1X TBST. After the last TBST wash, a final TBS wash was done. The membrane was then scanned with the side that was facing up (closest to gel during transfer) now facing down on the clean surface of a LI-COR Odyssey infrared fluorescence scanning instrument (LI-COR model number 9140).

Antibodies were obtained from and used with working concentrations as follows: p-MAPK (Cell Signaling, #4370S, 1:1,000), α-Tubulin (Novus, #NB100-690, 1:1,000), ß-actin (LI-COR #926-42212, 1:1,000), anti-rabbit (800CW LI-COR #926-32211, 1:20,000) anti-mouse (680LT LI-COR #925-68070, 1:20,000).

### Imaging and Quantification

Placement of the membrane on the scanning surface was set in Image Studio. Both 700nm and 800nm wavelength channels were set to be scanned. Resolution was set to 42 um, and focus offset was set to 0.0 mm. After the membrane finished scanning the image and associated zip file were exported from the Li-Cor Odyssey scanner and imported into Image Studio Lite for analysis. In Image Studio, boxes were drawn around protein bands to generate signal values.

## Acknowledgements

Funding for this project was in part provided by the NIH / NHGRI grant U54HG008098 and by
Clemson University. We thank Tim Pruett in the Clemson Additive Manufacturing core for 3D printing services, and Mark Ciaccio for helpful discussions.

